# Beacon-based sleep status and physical activity monitoring in humans

**DOI:** 10.1101/2023.10.30.564338

**Authors:** Takefumi Kikusui, Mizuho Yagisawa, Kahori Koyama, Koichi Fujiwara, Kazuhiko Kume, Kensaku Nomoto, Miho Nagasawa

**Affiliations:** Laboratory of Human-Animal Interaction and Reciprocity, School of Veterinary Medicine, Azabu University; Department of Material Process Engineering, Nagoya University, Nagoya, Aichi, 464-8601 Japan; Department of Neuropharmacology, Graduate School of Pharmaceutical Sciences, Nagoya City University, Nagoya, Aichi, 467-8603, Japan; Department of Physiology, Dokkyo Medical University School of Medicine, Mibu, Tochigi, 321-0293, Japan

**Keywords:** Beacon accelerator, Sleep-wake cycle, Exercise level, smartphone-monitoring

## Abstract

One out of every five people in Japan is dissatisfied with their sleep and various diseases caused by lack of exercise have been pointed out, but there are few effective remedies for these problems. In this study, we aimed to develop a simple method for measuring behavioral sleep patterns and physical activity using a beacon accelerometer wirelessly connected with a smartphone. A sleep prediction model was created comparing the data obtained from the accelerometer with the sleep status data obtained by a previously validated sleep monitoring system. The Random Forest model was able to classify sleep and wakefulness with a 97.4% and 85.4% precision, respectively, which were comparable to those of conventional acceleration-based sleep monitoring devices. Additionally, the same data acquisition method was used to classify exercise intensity into seven levels and a high correlation (r=0.813, p<0.0001) was found when comparing the classified exercise intensity to metabolic equivalent (MET) values. This suggests that the proposed method can be used for accurate measurement of both behavioral sleep and physical activity classifying over a long period of time.

## Introduction

Sleep quality and quantity are closely linked to both mental and physical health. According to a survey conducted by the Ministry of Health, Labor, and Welfare in Japan, one in five individuals report experiencing some form of sleep-related complaint, with this number increasing annually(1). Healthy adults typically experience a cycle of non-REM and REM sleep, each lasting approximately 1.5 hours, with several cycles comprising a night’s sleep. A deeper stage (N3) of non-REM sleep, typically occurs in the first two cycles, or about 3 hours after falling asleep. Waking up at the end of each cycle after falling asleep, is considered to contribute to a good quality sleep with a positive morning mood(2).

Various methods for objectively evaluating sleep quality have been proposed, and the polysomnography (PSG) is the most reliable method, but this method requires extensive equipment and is not practical for widespread use among the general population. Additionally, as nighttime sleep and daytime wakefulness are interconnected, it is important to evaluate both sleep quality and quantity, as well as daily physical activity(3). One method for this is the use of accelerometers, such as those found in smartwatches, to monitor sleep and activity. The head movement was monitored by accelerator and it can estimate the sleep stages with high accuracy(4). Algorithmic processing of data from accelerometers worn on the wrist enabled highly accurate classification of sleep and activity(5). However, a small proportion of these technologies have been independently validated and proven effective for research use (6,7). For example, Apple watch and Fitbit are widely used commercially but raw data was not accessible, and the determination of sleep and activity levels are not disclosed. Additional validation using various methods and comparisons will help establish appropriate measurement methods for sleep and activity monitoring for research purpose. Hence, there exists a requirement for additional validated techniques that are readily available. The current research endeavors to employ a commercially accessible 3-axis accelerometer to track both physical activity and sleep patterns and transmit the gathered data to a smartphone for analysis and validation purposes. In this study, the accelerometer was put on the west, not on the arm like smartphone, to monitor the precise whole body movement(8).

## Methods

In this study, the first objective was to develop a sleep prediction model utilizing machine learning techniques. Data obtained from body movement by an accelerometer was utilized as the training dataset. The model was trained using the sleep status data obtained by Omron Corporation’s HSL-102-M, a previously validated, clinically approved sleep monitoring system (9). The second objective was to assess the physical activity levels based on the same data acquisition methods, and the obtained data were utilized to assess the degree of concordance with the metabolic equivalent (MET) values.

## Experimental Materials

The following equipment were used. Bluetooth beacon accelerometer (MetamotinRL, Mbientlab Co. Ltd., CA, USA), Android smartphone (Aquos sense2, SHARP, Co, ltd, Tokyo, Japan), and Omron Sleep Meter HSL-102-M (Tokyo, Japan). The accelerometer, in the form of a beacon, was worn on the waist to measure 3-axis acceleration (X-, Y- and Z-axis). The beacon and the smartphone were connected via BLE (Bluetooth Low Energy) to transmit the acceleration data. The detection range of the beacon was approximately 10 meters, and the smartphone was placed in a static location within this range when the user was at home or sleeping, and carried with them when they were out. An application for data acquisition was developed in collaboration with Quadlytics Inc (Kyoto, Japan). The data acquired included timestamp, username, beacon identifier, 3-axis acceleration, radio wave strength (rssi), latitude, and longitude, and the sampling rate was 12.5 Hz.

The sleep monitor Omron Sleep Meter used 10.5 GHz radio waves emitted from the main unit to capture tossing and turning and chest movements during sleep, and measures the time spent in the sleep/wake state based on the frequency and magnitude of body movements. The data recorded on the sleep meter was extracted using Omron’s Sleep Design Viewer, a dedicated data management software, via SD card. Sleep data were classified into four sleep statuses determined by the duration of body movements every 30 seconds: status 1 = asleep, status 2 = awake, status 3 = absent, and status 4 = unknown. Sleep and awake data were used as teaching data and those classified as other or absent were not used.

## Subjects

The experimental procedures were approved by the Ethical Committee for Medical and Health Research Involving Human Subjects of Azabu University (#052). Eight participants (one male and 7 females, age 19-32 (average 23.7, SD 4.4)) were included in the sleep data analysis and 17 participants (6 males and 11 female, age 21-70 (average 47.0, SD 15.3)) were included in the activity categorizing analysis. All of them had no clinical symptoms of sleep disorder or physical disabilities. They were informed about the procedure of the experiments, and all of them gave an informed consent before taking part in the experiment. All subjects were living a regular life and there was no restriction from caffeine intake, drinking and excessive exercise, due to monitor a daily life in ordinary people.

## Procedure

A sleep monitoring device and a smartphone were placed in a fixed location within the subject’s bedroom to ensure proper detection range. A beacon device, which was attached to the subject’s waist, was utilized to record data on the subject’s sleep patterns. The subjects initiated the measurement process by pressing the “start” button on the sleep monitoring device before going to bed and pressing the “end” button upon waking. To ensure that only sleep data was analyzed, the beacon batteries were inserted prior to bedtime and removed upon waking. The experiment was conducted over a period of seven days and any beacon data with a high number of missing values due to unstable Bluetooth connections were excluded from the analysis and results (22.3% of total data).

## Data analysis

Data from an acceleration sensor were processed using the MATLAB programming code (MathWorks, Co. Ltd, Natick, MA, USA). The data from the three axes of the sensor were combined into a scalar value and then averaged over a one-second interval. As the Omron Sleep Meter, the device used for measuring sleep, exported data in 30-second epochs, the standard deviation of the 30-second acceleration data was calculated and used as the data for the 30-second epochs.

As previously described the monitoring sleep-wake cycle by accelerator, SD (standard deviation) values of vector quantities in the three axes converted to scalar values of acceleration were calculated every 30 seconds. Machine learning was then performed using five consecutive variables including one minute before (2 data) and one minute after (2 data) the target epoch (10). These data were then grouped as a single unit and matched to the sleep monitor Omron Sleep Meter training data (status 1 and 2). Machine learning algorithms including Random Tree, Random Forest and J48 were applied to classify the data. The parameters are determined using Bayesian optimization. The trees are pruned to the extent that the error is not increased by 10 times of cross-validation.

Similarly, the diurnal activity data of a time series of 30-second of SD data samples, which were collected for one minute prior to and one minute following 2 designated epochs, were subsequently analyzed. Normal mixed clustering was performed, and the activity levels were classified into seven clusters (JMP, v13.0, SAS Institute. Ltd, Cary, NC, USA). The number of activity clusters (low to high) followed the MET scale, where a value of 1 corresponds to resting metabolism and a value of approximately 7 corresponds to the highest level of daily physical activity. Higher numbers in the clusters indicate higher activity. Based on this teacher data, automatic classification of activity levels was carried out using MATLAB random tree protocol with an optimization as described above.

Finally, correlation analysis between the automated-classified activity data, and metabolic equivalent (MET) values, which is commonly used to evaluate the intensity of daily physical activity(11). As the accelerometer-based exercise measurement utilized in this study was not designed to capture high-intensity physical activities such as those performed in competitive sports, MET was validated for intensities up to a maximum of 7. The participants were asked to perform 3 minutes of activity at the exercise intensity described in the MET values and the data was used for comparative analysis.

## Results

### Accuracy of prediction of sleep status

Figure 1A presents an instance of 1-second acceleration data, and the SD of the 30-second acceleration data during daytime. Figure 1B displays the SD values obtained from the accelerometer data and body movement data obtained by Omron Sleep Meter, during the nighttime. The data was categorized using machine learning algorithms, including Random Tree, Random Forest, and J48, and cross-validation (10% of cases) results were provided in Table 1. The prediction of sleep status 1 (asleep) was comparable across all algorithms (97.4-98.1%). However, the prediction accuracy of sleep status 2 (awake) was observed to be higher by using the Random Forest algorithm (85.4%) as compared to the other two algorithms (Table S1). F1 scores were also calculated and it was revealed that F1 scores in Status 1 was comparable among the three algorithms (0.984-0.987) but Random Tree showed high F1 score in Status 2 (Table S1, 0.447).

**Figure 1.**
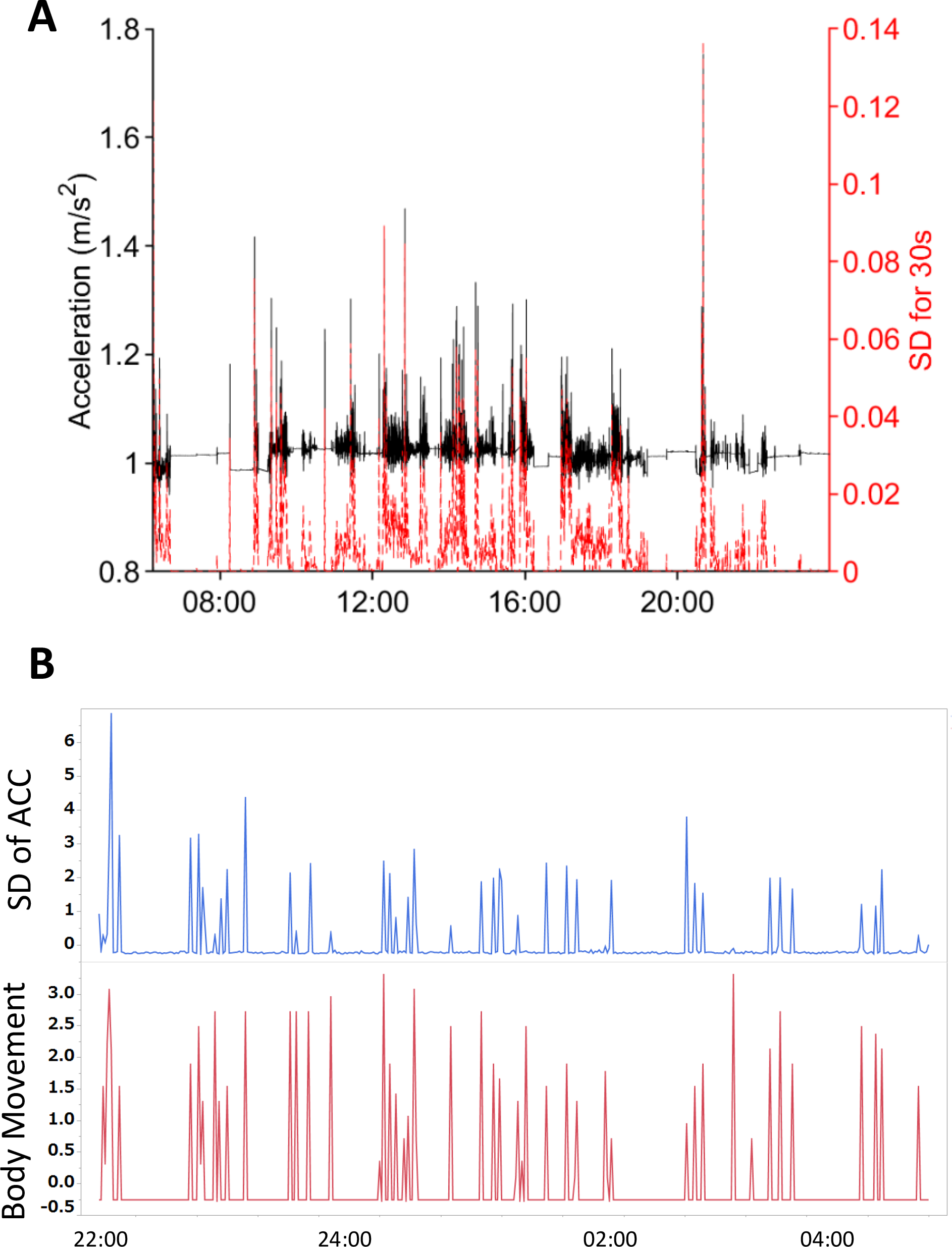
(A) An example of the raw data of vector quantities in the three axes converted to scalar values of acceleration obtained by the beacon (black line and the first Y axis) and the data SD values of them calculated every 30 seconds (red line and the second Y axis). X axis represented the time of the day. (B) A representative data of the SD values of acceleration obtained by the beacon (upper, blue line) and body movement data obtained by Omron Sleep Meter (lower, red line).

**Table 1.** Results of machine learning predictive values of sleep stages.

| Algorism | Sleep Status |  |
| --- | --- | --- |
|  | 1 | 2 |
| Random Forest | 0.974 | 0.854 |
| Random Tree | 0.981 | 0.482 |
| J48 | 0.975 | 0.621 |
| Sleep status1: asleep |  |  |
| Sleep status2: awake |  |  |

### Physical activity classification

The SD values derived from the similarly obtained accelerometer data were analyzed using Normal mixed clustering to categorize activity levels into seven distinct stages. The Mean SD values of each of the seven activity stages were compared, as illustrated in Figure 2A. These results were then employed in a Random Forest method, and the 10 times of cross-validation results yielded an accuracy of 97.0% for 10% of the cases.

**Figure 2.**
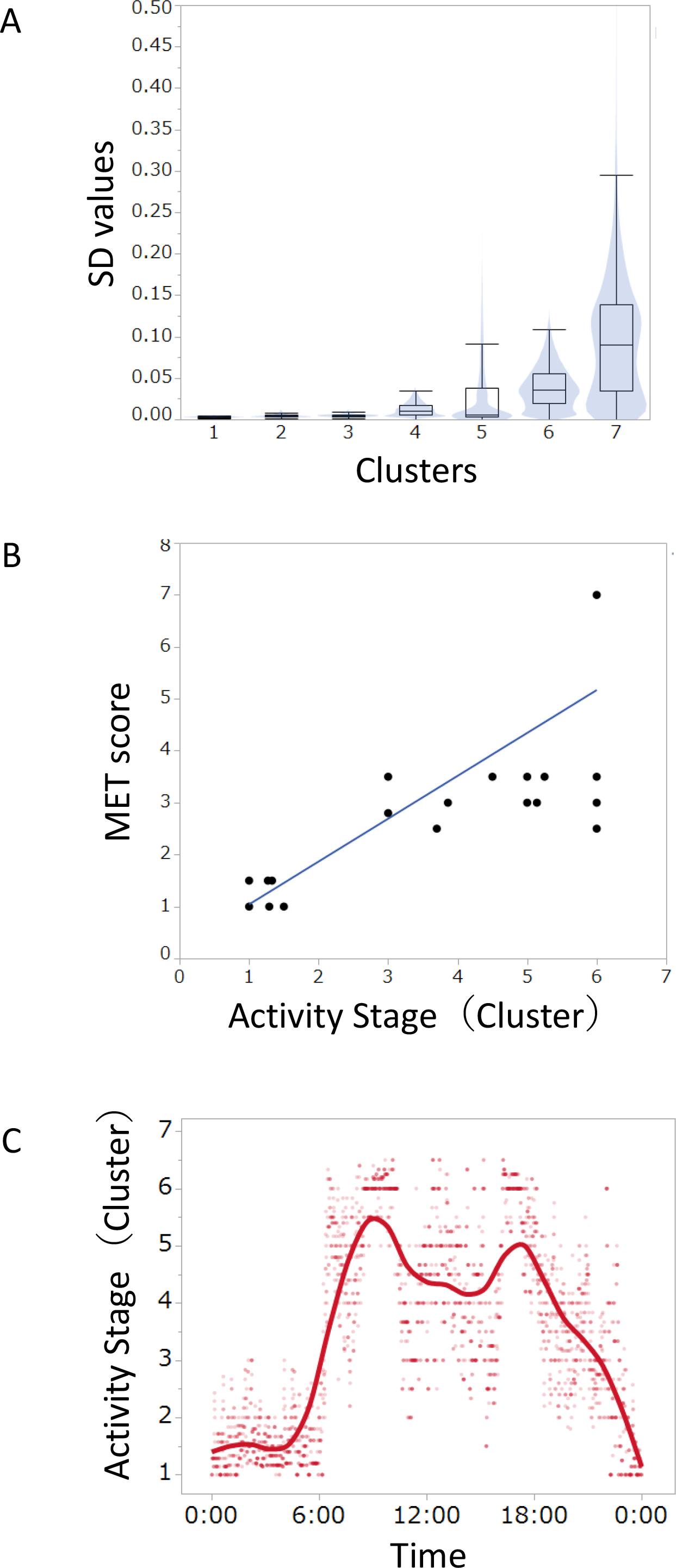
(A) The mean SD values of each of the seven activity stages. (B) Correlation analysis of activity stages (7 clusters) and the MET score (r=0.813, p<0.0001). (C) A representative data of daytime and nighttime activity cycles classified by machine learning. Note that there was a clear day-night activity change.

Figure 2B represents a comparison between activity stages and the MET score, which revealed a significant correlation between the two (r=0.813, p<0.0001).

Lastly, data was collected continuously from adult males for seven days, and machine learning techniques were used to categorize activity stages into seven clusters. The collected data was plotted from midnight to 24:00 hours, and a smoothing spline was fitted with a parameter value of λ=0.25. The resulting analysis illustrated distinct daytime and nighttime activity cycles, as demonstrated in Figure 2C.

## Discussion

The current investigation employed machine learning algorithms to analyze data collected from accelerometers, which yielded results that closely corresponded to those obtained from the validated sleep meter. This suggests that the data processing technique employed in this study is valuable for detecting sleep status. Sleep status 1 (asleep) was classified with greater than 97% accuracy, whereas status 2 (awake) was classified with a lower accuracy of 85% by random tree classification. This lower accuracy may be attributed to limitations in detecting body movement with the accelerometer, which was attached to the waist and thus unable to detect movements of the arms and head. In contrast, sleep meters employ whole-body sensing using radio-wave radiation, which can detect small movements. An alternative method for sleep monitoring is to use smartwatches attached to the wrist, which can detect arm movement. In this study, an Omron sleep monitor was used for machine learning of sleep classification by accelerometer. Although PSG-based validation was ideally needed, a Omron sleep monitor was chosen due to the difficulty of the PSG experiment. In the future, it will be necessary to verify the degree to which the algorithm used in this study agrees with PSG results. The accelerometers were placed on the waist to measure whole-body movement as an indicator of activity. The choice to avoid using an accelerometer on the arm was made due to concerns that it would generate high values when the hands are engaged in tasks. Nonetheless, measuring sleep with an accelerometer on the arm represents a future challenge.

The method of analyzing the data together with the pre- and post- data is in accordance with previous research on sleep assessment using accelerometers(10,12). In a previous study, an algorithm was developed and tested for effectiveness in assessing sleep/wake every 2 minutes from 4 minutes before to 4 minutes after the relevant time of activity intensity measured by a waist-mounted accelerometer. The results showed that the average agreement with the corresponding PSG-based sleep/wake data was 86.9%, the average sensitivity (sleep detection) was 89.4% and the average specificity (wake detection) was 58.2%(12). These accuracies were corresponding to the present study, suggesting that 1 minute pre- and 1 minute post- data are sufficient for the detection of sleep status.

Although the classification of sleep stages could not be reached in this analysis, if this analysis becomes possible, it would make a significant contribution to human medical research. This is because different functions have been reported for each sleep stage. In humans, non-REM sleep appears at the onset of sleep, followed by REM sleep one to two hours later. This cycle is repeated four to five times during the night. During non-REM sleep, sympathetic nervous system activity rests and breathing and heart rate calm down(2). Characteristic brain waves called slow waves are observed in the cerebral cortex, and several physiological roles have been identified, including growth hormone secretion and removal of metabolites accumulated in the brain(13). On the other hand, during REM sleep, rapid eye movement (REM), from which the name is derived, occurs, accompanied by a loss of skeletal muscle tone and a decrease in thermoregulation. Sympathetic nerve activity increases to the same level or higher than during wakefulness(14). REM sleep is thought to contribute to memory consolidation by being involved in the generation of two types of brain activity: slow waves and theta waves (13), and recent studied demonstrated that non-REM N2 sleep is also contributed to memory consolidation (15).

In this study, a comparable data processing approach was employed to classify physical activity. When standard deviation (SD) values were categorized into seven clusters, these clusters exhibited strong correlations with Metabolic Equivalent of Task (MET) scores, indicating the potential for accurate and easy-automated evaluation of physical movement. MET is a widely utilized physiological concept that provides a simple means of quantifying energy expenditure during physical activities. For instance, elderly Japanese men displayed a correlation between health status and daily activity exceeding 3 MET(11). In the present study, MET values were estimated from the data obtained using an accelerometer, which demonstrates the usefulness of automatic measurement of physical activity via accelerometers for the assessment of daily exercise and its impact on physical and mental health(11).

The data processing in the present study was performed as a post-hoc analysis in MATLAB, but the process was not overly complicated. Consequently, the sensor data can be transmitted to a smartphone and calculated on the device. In the future, sleep and activity levels may be displayed on smartphones, leading to personalized recommendations for sleep and activity levels.

## Acknowledgements

The authors thank Dr. Yuya Hataji for his kind help for the MATLAB coding.

## Reference list

1. Tanaka H, Shirakawa S. Sleep health, lifestyle and mental health in the Japanese elderly: ensuring sleep to promote a healthy brain and mind. J Psychosom Res. 2004 May;56(5):465–77.

2. Gradisar M, Gardner G, Dohnt H. Recent worldwide sleep patterns and problems during adolescence: a review and meta-analysis of age, region, and sleep. Sleep Med. 2011 Feb;12(2):110–8.

3. Vallat R, Berry SE, Tsereteli N, Capdevila J, Al Khatib H, Valdes AM, et al. How people wake up is associated with previous night’s sleep together with physical activity and food intake [Internet]. Vol. 13, Nature Communications. 2022. Available from: 10.1038/s41467-022-34503-2

4. Yoshihi M, Okada S, Wang T, Kitajima T, Makikawa M. Estimating Sleep Stages Using a Head Acceleration Sensor. Sensors [Internet]. 2021 Feb 1;21(3). Available from: 10.3390/s21030952

5. Ode KL, Shi S, Katori M, Mitsui K, Takanashi S, Oguchi R, et al. A jerk-based algorithm ACCEL for the accurate classification of sleep-wake states from arm acceleration. iScience. 2022 Feb 18;25(2):103727.

6. Peake JM, Kerr G, Sullivan JP. A Critical Review of Consumer Wearables, Mobile Applications, and Equipment for Providing Biofeedback, Monitoring Stress, and Sleep in Physically Active Populations. Front Physiol. 2018 Jun 28;9:743.

7. Sadeh A. The role and validity of actigraphy in sleep medicine: an update. Sleep Med Rev. 2011 Aug;15(4):259–67.

8. Fanning J, Miller ME, Chen S-H, Davids C, Kershner K, Rejeski WJ. Is Wrist Accelerometry Suitable for Threshold Scoring? A Comparison of Hip-Worn and Wrist-Worn ActiGraph Data in Low-Active Older Adults With Obesity. J Gerontol A Biol Sci Med Sci. 2022 Dec 29;77(12):2429–34.

9. Hashizaki M, Nakajima H, Tsutsumi M, Shiga T, Chiba S, Yagi T, et al. Accuracy validation of sleep measurements by a contactless biomotion sensor on subjects with suspected sleep apnea. Sleep Biol Rhythms. 2014 Apr 1;12(2):106–15.

10. Cole RJ, Kripke DF, Gruen W, Mullaney DJ, Gillin JC. Automatic sleep/wake identification from wrist activity. Sleep. 1992 Oct;15(5):461–9.

11. Aoyagi Y, Shephard RJ. Habitual physical activity and health in the elderly: the Nakanojo Study. Geriatr Gerontol Int. 2010 Jul;10 Suppl 1:S236–43.

12. Enomoto M, Endo T, Suenaga K, Miura N, Nakano Y, Kohtoh S, et al. Newly developed waist actigraphy and its sleep/wake scoring algorithm [Internet]. Vol. 7, Sleep and Biological Rhythms. 2009. p. 17–22. Available from: 10.1111/j.1479-8425.2008.00377.x

13. Stich FM, Huwiler S, D’Hulst G, Lustenberger C. The Potential Role of Sleep in Promoting a Healthy Body Composition: Underlying Mechanisms Determining Muscle, Fat, and Bone Mass and Their Association with Sleep. Neuroendocrinology. 2022;112(7):673–701.

14. Somers VK, Dyken ME, Mark AL, Abboud FM. Sympathetic-nerve activity during sleep in normal subjects. N Engl J Med. 1993 Feb 4;328(5):303–7.

15. Schönauer M. Sleep Spindles: Timed for Memory Consolidation. Curr Biol1 Elsevier; 2018 Jun 4;28(11):R656–8.

