## Supplemental Table 1 for "Beacon-based sleep status and physical activity monitoring in humans"

**Raw data points**

**Random Tree**

| St1 | St2 | | classified |
|---|---|---|---|
| 21674 | 309 | 21983 | St1 |
| 404 | 288 | 692 | St2 |

status 1: 0.981. status 2: 0.482

**Random Forest**

| St1 | St2 | | classified |
|---|---|---|---|
| 21964 | 25 | 21989 | St1 |
| 547 | 147 | 694 | St2 |

status 1: 0.974. status 2: 0.854

**J48**

| St1 | St2 | | classified |
|---|---|---|---|
| 22735 | 45 | 22780 | St1 |
| 576 | 74 | 650 | St2 |

status 1: 0.975. status 2: 0.621

**Predictive Value**

**Random Tree**

| Precision | Recall | Accuracy | F1 Score |
|---|---|---|---|
| 0.982 | 0.986 | 0.969 | 0.984 |
| 0.482 | 0.416 | | 0.447 |

**Random Forest**

| Precision | Recall | Accuracy | F1 Score |
|---|---|---|---|
| 0.976 | 0.999 | 0.975 | 0.987 |
| 0.855 | 0.212 | | 0.339 |

**J48**

| Precision | Recall | Accuracy | F1 Score |
|---|---|---|---|
| 0.975 | 0.998 | 0.973 | 0.987 |
| 0.622 | 0.114 | | 0.192 |
